# Metatranscriptome data support the existence of two distinct morphotypes in a single parmalean species in natural environments

**DOI:** 10.64898/2026.04.16.718586

**Authors:** Hiroto Sasaki, Hisashi Endo, Eric Pelletier, Shiya Yoshikawa, Akira Kuwata, Hiroyuki Ogata

## Abstract

Parmales (Bolidophyceae), the closest relatives of diatoms, includes isolates exhibiting one of two distinct morphotypes: a silicified non-flagellated form (S-type) and a naked flagellated form (F-type). Although alternation of these two forms for a single isolate has not been formally established, previous studies hypothesized that these morphotypes represent alternating stages of a single parmalean. In this study, we investigated the global expression patterns of S- and F-type marker genes by integrating parmalean Metagenome-Assembled Genomes (MAGs) and the metatranscriptomic dataset from *Tara* Oceans. We detected the expression of both S- and F-type marker genes from individual environmental MAGs. This finding provides the first metatranscriptomic evidence that natural parmalean genomes possess the potential to manifest both morphotypes. Furthermore, our analysis revealed different geographical expression patterns between the two forms. The expression of F-type markers showed a broad distribution, whereas that of S-type markers was more restricted, suggesting distinct niches for the two morpho-phases. Moreover, S-type gene expression appears to require specific environmental triggers that lead to a higher population density, whereas F-type expression is rather constitutively maintained. Overall, our results support the hypothesis of a life cycle involving morphological switching and reconcile the long-standing discrepancy between the ubiquity of parmaleans in molecular surveys and the limited geographical range for the observation of silicified cells. Based on these patterns, we propose a threshold-based model in which F-type-dominated populations persist under conditions unfavorable for growth and a morphological switch to the S-type is triggered once environmental conditions exceed a critical threshold for growth.

## Introduction

Diatoms are one of the most ecologically successful groups of phytoplankton, accounting for approximately 20% of primary production in modern oceans (Armbrust, 2009). However, the drivers of their ecological success remain elusive, and comparative studies involving their close relatives are increasingly recognized as promising approaches to address this question (Ichinomiya et al., 2016).

The order Parmales comprises pico-eukaryotic algae that form the sister group of diatoms (Guillou et al., 1999; Ichinomiya et al., 2011). Originally, two morphological forms have been described: a silicified form surrounded by silica plates (S-type) (Booth & Marchant, 1987), and a naked flagellated form possessing two unequal flagella (F-type) (Guillou et al., 1999). Recently, a novel silicified morphotype covered with distinct silica scales, referred to as ‘Scaly parma’, was identified within the basal lineage of Parmales (Ban et al., 2023). This discovery led to the recent establishment of the order Lepidoparmales to accommodate this lineage (Kamakura et al., 2025). Although these two morphological forms were historically regarded as distinct lineages, molecular phylogenetic analyses revealed their close relationship (Ichinomiya et al., 2011). Consequently, both S- and F-type algae— originally classified separately—were reclassified into the single order Parmales within the class Bolidophyceae (2). Ecologically, S-type cells are most abundant in polar and subarctic regions (Booth et al., 1980; Ichinomiya & Kuwata, 2015), whereas F-type populations were originally isolated from the equatorial Pacific Ocean and the Mediterranean Sea (Guillou et al., 1999). Despite these habitat preferences based on isolation, microscopic observations and molecular studies indicate that the algae in Bolidophyceae are distributed globally (Ban et al., 2024; Silver et al., 1980) and the morphological forms display different distributions across depth and seasons (Kuwata et al., 2018). These algae are referred to as parmaleans in this study.

The close evolutionary relationship between the S-type and F-type parmaleans raised the possibility that these morphotypes correspond to different life stages of a single organism. Mann and Marchant hypothesized that the life cycle of ancestral diatoms consisted of haploid flagellated cells and diploid silicified zygotes akin to that of the early branching centric diatoms (Mann & Marchant, 1989). This type of life cycle is also known for other algae such as the haptophyte *Gephyrocapsa huxleyi* (Green et al., 1996). Given the evolutionary relatedness between S-type and F-type parmalean strains and their sister relationship to diatoms, Ichinomiya et al. (2011) hypothesized that Parmales possess a life cycle that switches between a diploid silicified form and a haploid flagellated form (Ichinomiya et al., 2011). This hypothesis was supported by several lines of evidence. First, a study of a flagellated strain revealed a low level of heterozygosity, suggesting the haploid status of the genome for this strain (Kessenich et al., 2014). Second, the flagellated strain *Triparma* sp. RCC1657 was found to belong to a clade of silicified forms based on SSU rRNA gene sequences, despite its morphology (Ichinomiya et al., 2016). Third, a genome sequencing study of silicified strains revealed the presence of genes related to the flagellated form in their genomes led to the hypothesis that the hypothesized life cycle switching corresponds to an alternation between a silicified photoautotrophic stage and a naked flagellated phago-mixotrophic stage (Ban et al., 2023). Specifically, Ban et al. (2024) identified genes associated with phagocytosis in parmalean genomes—genes that have been lost in diatoms—and proposed that this mixotrophic strategy facilitates their adaptation to varying nutrient environments. Furthermore, a recent global metabarcoding study consistently revealed cosmopolitan distributions of parmaleans, which may be explained by their life cycle strategy (Ban et al., 2024). However, to our knowledge, the life cycle switches between the two morphological forms in a single parmalean has not been formally established, although Ichinomiya et al. (2016) mentioned the appearance of flagellated cells in cultures of silicified parmaleans (Ichinomiya et al., 2016). Therefore, the hypothesis on the life cycle and genome state of parmaleans remains to be tested.

To assess the hypothesis, large-scale environmental omics resources provide a powerful framework. The *Tara* Oceans expedition generated global-scale metagenomic and metatranscriptomic datasets for surface and mesopelagic plankton (Sunagawa et al., 2020). Among its key outputs, the Marine Atlas of *Tara* Oceans Unigenes (MATOU-v1.5) provides a comprehensive catalog of eukaryotic gene expression in the marine environment (Carradec et al., 2018; Pierella Karlusich et al., 2025).

In this study, we examine the hypothesis that the S-type and F-type forms represent distinct life-cycle stages of a single parmalean. Under this hypothesis, genes specific to S-type or F-type forms would be expressed from a single parmalean, possibly in different geographic locations or environmental conditions. Specifically, we aim to clarify whether the genes related to silicified or flagellated forms are expressed from individual parmalean populations (at the species level) in the ocean and whether the geographic distributions of these gene expressions are environmentally regulated. To achieve this, we first select marker genes representing the two distinct morphological forms based on differential gene expression between cultured strains based on publicly available transcriptome data. We then examine the presence of these genes in two parmalean Metagenome-Assembled Genomes (MAGs) derived from *Tara* Oceans metagenomic data (Delmont et al., 2022). By integrating these MAGs with the *Tara* Oceans metagenome and MATOU dataset, we characterize the global expression patterns of these marker genes and correlate them with the environmental parameters.

## Materials and Methods

### Transcriptome data

We analyzed publicly available RNA-seq data from S-type and F-type parmalean strains. For the S-type strain, raw reads of *Triparma laevis* f. *inornata* strain NIES-2565 [six samples: three silicate-rich (concentration of Si, 50 μM) and three silicate-deficient (0.23 μM) replicates] were retrieved from NCBI Sequence Read Archive (SRA) (BioProject PRJDB4267). Both conditions were treated as S-type in this study because the overall expression profiles were quiasi-indistinguishable between conditions (see **Fig. S1**), though a previous study reported that the strain tend to lose their silica plates under silicate-deficient conditions (Yamada et al., 2014). For the F-type strains, data for four strains (*Triparma eleuthera* CCMP1866, *Triparma* sp. RCC1657, RCC208, and RCC2347) were obtained from the MMETSP database (Keeling et al., 2014).

Reads were quality-filtered with fastp v0.24.0 (Chen et al., 2018), rRNA-depleted with SortMeRNA v4.3.7 (Kopylova et al., 2012) using the PR2 database v5.1.0 (Guillou et al., 2013), assembled de novo with Trinity v2.15.2 (Grabherr et al., 2011), and redundancy-reduced using the EvidentialGene pipeline (https://sourceforge.net/projects/evidentialgene/) to obtain non-redundant CDS and protein sequences for each strain.

### Gene expression quantification and completeness assessment

To quantify gene expression levels, the quality-filtered RNA-seq reads were mapped back to the constructed non-redundant reference gene catalogs using Salmon (version 1.10.3) (Patro et al. 2017) in quasi-mapping mode with default parameters. This provided transcript abundance estimates for each sample, which were subsequently used for downstream differential expression analysis.

To assess the completeness of the assembled transcriptomes, we performed BUSCO v5.8.3 analysis (Manni et al., 2021). The non-redundant predicted protein sequences from the five transcriptomes were evaluated against the stramenopile_odb12 lineage dataset. For comparative purposes, the same BUSCO assessment was also performed on the coding sequences of eight publicly available parmalean genomes.

### Orthology inference

To investigate evolutionary relationships and identify core gene sets, we performed an orthology analysis using OrthoFinder v2.3.8 (Emms & Kelly, 2019). The input dataset included the predicted protein sequences from (1) the five assembled transcriptomes (one S-type and four F-types described above), (2) eight publicly available parmalean genomes, (3) two parmalean Metagenome-Assembled Genomes (MAGs) retrieved from the *Tara* Oceans dataset: TARA_ARC_108_MAG_00221 (ARC-MAG) and TARA_PON_109_MAG_00217 (PON-MAG), and (4) three diatom genomes (*Thalassiosira pseudonana, Pseudo-nitzschia multistriata, Fragilariopsis cylindrus*) included as outgroups.

### Phylogenetic analysis

Based on the OrthoFinder results, single-copy orthologous genes conserved across all genomes and transcriptomes were extracted to reconstruct the phylogeny. The sequences were concatenated and aligned, and a maximum likelihood phylogenetic tree was constructed using IQ-TREE 3.0.1 (Wong et al., 2025). ModelFinder (Kalyaanamoorthy et al., 2017) was used to determine the best-fit substitution model, and branch support was assessed using ultrafast bootstrapping (Hoang et al., 2018).

### Identification of morphotype-specific marker genes

From the OrthoFinder output, we extracted orthogroups conserved across all eight Parmales reference genomes and filtered for genes with high expression levels (logCPM (Counts per million) > 5). Marker genes were then classified based on their exclusive expression in either S-type or F-type cultures. This conservative classification strategy was adopted to minimize false positives arising from inter-strain differences.

### Metagenomic and metatranscriptomic mapping to MAGs

Morphotype-specific marker gene sequences were extracted from the two parmalean MAGs (ARC-MAG and PON-MAG) based on orthogroup assignments. Metagenomic and metatranscriptomic reads were mapped to these MAG-derived marker genes using bwa-mem2 v2.2.1 (Vasimuddin et al., 2019) with default parameters. The resulting alignments were sorted and deduplicated using samtools v1.18 (Danecek et al., 2021). To ensure high-confidence mapping (Carradec et al., 2018), reads were filtered using bamFilters v1.10.2 (https://github.com/institut-de-genomique/bamFilters) with a minimum sequence identity of 95% and a minimum alignment length coverage of 90%. Finally, mapped reads were extracted, and raw read counts were aggregated per sample to construct a count matrix.

### Clustering and visualization of expression patterns

A Bray-Curtis dissimilarity matrix was calculated from the marker gene expression matrix using the vegan package v2.7-1 in R v4.5.1, and hierarchical clustering was performed using the Ward.D2 method. Results were visualized as heatmaps with clustering dendrograms using the pheatmap package v1.0.13. Expression data were also integrated with *Tara* Oceans environmental metadata.

### Environmental modeling

To investigate the relationships between morphotype expression and environmental factors, we employed Generalized Additive Models (GAMs) using the mgcv package v1.9.3. In this modeling framework, the response variable was defined as a normalized expression index calculated using log10-transformed reads with a pseudo count of 1. Specifically, the index for each type (F or S) was calculated as:

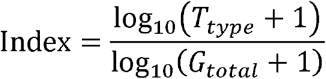

where *T*_*type*_ represents metatranscriptomic read counts for type-specific marker genes (F- or S-type), and *G*_*total*_ represents the sum of metagenomic read counts for both F- and S-type genes. Samples with low gene abundance (*G*_*total*_ < 10) were excluded from the analysis. The explanatory variables included nutrient concentrations (total carbon, PO_4_, Si, and NO_2_+NO_3_) and physical parameters (temperature, salinity, and density). Sampling depth (Surface [SRF] or Deep Chlorophyll Maximum [DCM]) was included as a categorical factor. Prior to analysis, we confirmed that the Variance Inflation Factor (VIF) for all continuous predictors was less than 10. The model assumed a Tweedie error distribution with a log link function, and parameter estimation was performed using REML with the “select = TRUE” option enabled. This applies additional shrinkage, allowing the model to shrink non-contributing smooth terms toward zero, effectively removing them.

## Results

### De novo transcriptome assembly of five cultured Parmales strains

To explore possible morphotype-specific genes in parmaleans, we generated de novo RNA-seq assembly for five strains: the silicified strain NIES-2565 (n=6) and four flagellated strains (CCMP1866, RCC1657, RCC208, and RCC2347; n=1 each), yielding 19,000–30,000 transcripts per strain (**(6)**). BUSCO analysis (stramenopiles_odb12) confirmed completeness scores exceeding 60% for all assemblies, which is comparable to the reference complete genomes.

### Identification of morphology-specific markers

Prior to marker gene selection, we assessed whether the silicate-rich and silicate-deficient culture conditions of NIES-2565 produced distinct expression profiles. A Principal Coordinates Analysis (PCoA) based on Bray-Curtis dissimilarity of ortholog-mapped read counts across all ten samples revealed that the first principal coordinate (PCoA1), accounting for 63.3% of the total variance, clearly separated the four F-type strains from all six NIES-2565 samples. The silicate-rich and silicate-deficient replicates of NIES-2565 were intermingled along this axis, indicating that culture condition had a negligible effect on global expression patterns relative to the morphotype difference. Therefore, all six NIES-2565 samples were treated as S-type replicates in subsequent marker gene selection.

To identify genes specific to S-type or F-type forms, we conducted an ortholog analysis of 18 strains (five transcriptomes, eight parmalean genomes, and five diatom outgroups) and yielded 25,752 orthogroups. Of 5,638 orthogroups conserved across all eight reference genomes (“Parmales core gene set”), 39 were expressed exclusively in silicified cultures and 652 exclusively in flagellated cultures. Applying a stringent expression threshold (logCPM > 5) resulted in 30 S-type and 180 F-type marker genes (**Fig. 1**). Functional annotation confirmed that S-type markers included the silicon transporter (SIT) and F-type markers contained intraflagellar transport (IFT) genes, validating their correspondence to the silica cell wall and flagella, respectively. ARC-MAG and PON-MAG, the two parmalean MAGs that we analyze in this study, encoded 92% (166/181) of F-type and 73% (22/30) of S-type markers, while PON-MAG encoded 81% (139/181) and 90% (26/30), respectively.

**Figure 1.**
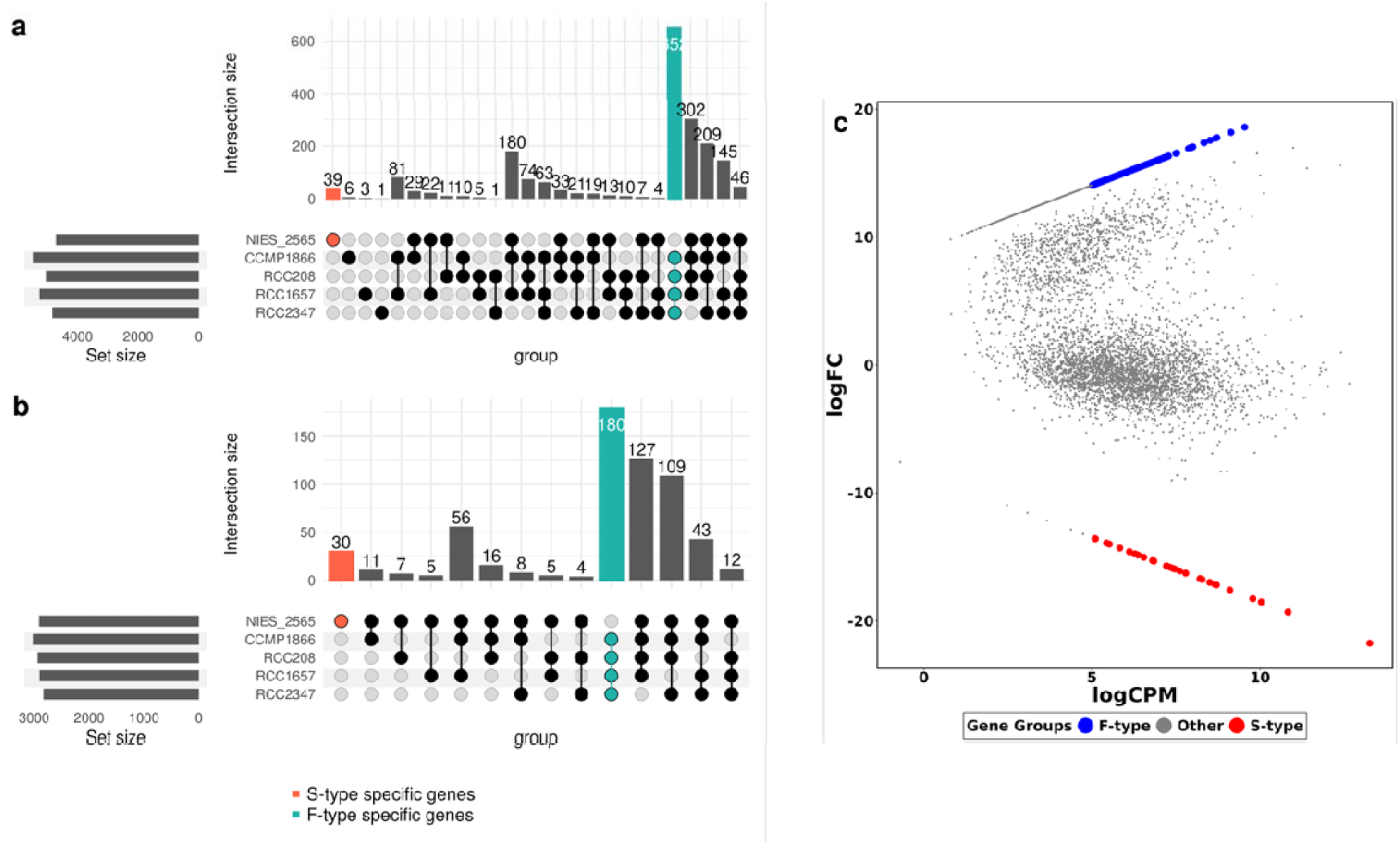
Genes specific to Parmales morphotypes. (a, b) UpSet plots illustrating the distribution of orthologs among different Parmales strains with transcriptome data. The numbers above the vertical bars (Intersection Size) indicate the number of orthologs shared by a specific combination of strains. The horizontal bars on the left (Set Size) show the total number of orthologs within each strain. The red bar corresponds to the orthologs unique to the S-type transcriptome, while the teal bar represents the orthologs identified in all F-type transcriptomes but not in the S-type. Panel (a) includes all orthologs that were found to be universally conserved in eight Parmales strain with genomic sequence data. In panel (b), only orthologs with an expression level of logCPM ≥ 5.0 are shown. (c) MA plot showing genes specifically expressed in individual morphotypes. Red points indicate orthologs selected as S-type specific and blue ones indicate orthologs selected as F-type specific.

### Phylogenetic relationships and MAG placement

To ensure our MAGs were representatives of the Parrmales known diversity, we inferred A phylogenetic tree based on 75 concatenated single-copy orthologous genes, which reproduced the established classification of Parmales with full bootstrap support (**Fig. 2**). The two MAGs were placed in distinct clades: ARC-MAG in the Lepidoparmales (Clade I) and PON-MAG in the *Tetraparma* group (Clade IV), representing two phylogenetically distant lineages. Average Nucleotide Identity (ANI) analysis further confirmed the ARC-MAG’s close affinity to the *Lepidoparma frigida* reference genome (99.4% identity).

**Figure 2.**
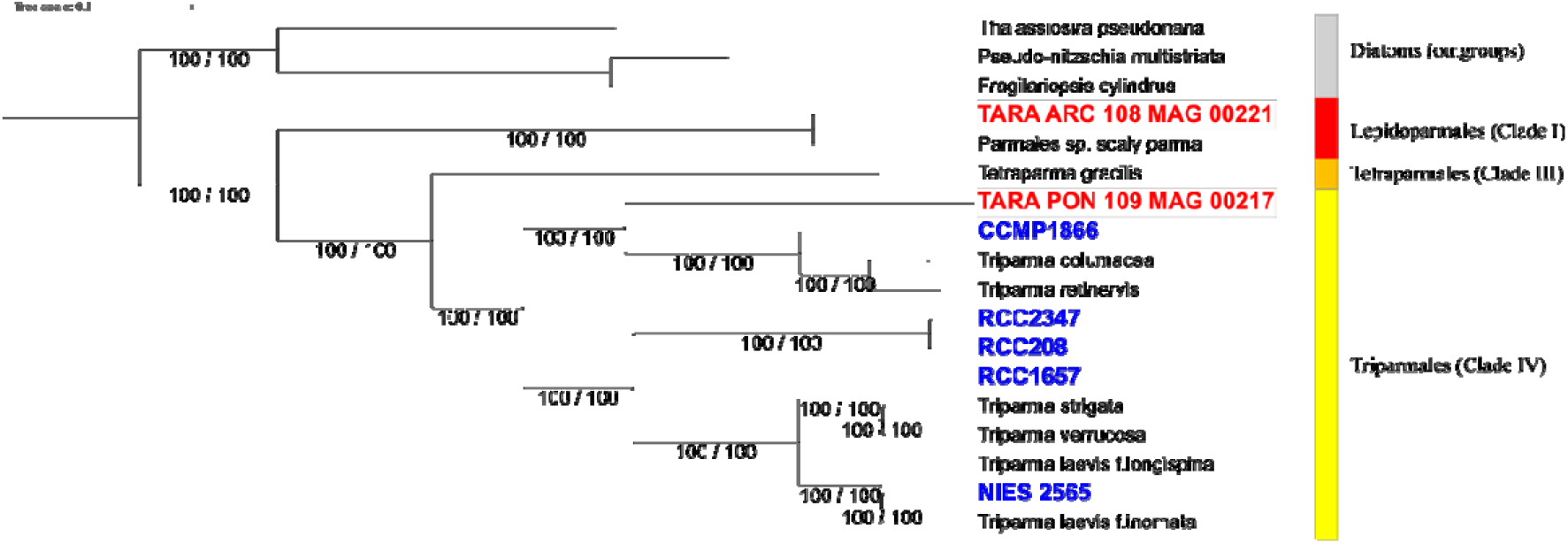
Phylogenetic relationships inferred from concatenated single-copy genes. Maximum likelihood phylogenetic tree of concatenated alignments of single-copy orthologous genes. IDs shown in blue represent the phylogenetic placement of assemblies derived from RNA-seq data. IDs shown in red indicate the positions of parmalean metagenome-assembled genomes (MAGs) from the *Tara* Oceans dataset. Nodal support is provided by 1,000 ultrafast bootstrap replicates (UFB, first value) and SH-aLRT (SH, second value).

### In situ expression of both morphotype markers from individual MAGs

To investigate the geographic distributions of S-type and F-type markers, we mapped *Tara* Oceans metagenomic and metatranscriptomic reads to the two MAGs. Metagenomic abundances (i.e., “presence”) of S-type and F-type markers were highly correlated within each MAG, confirming stable genomic copy number ratios (**Fig. S2, Table 2**). In contrast, metatranscriptomic abundances (i.e., “activity”) varied substantially across samples, suggesting that marker expression is dynamically regulated by local environmental conditions.

**Table 1.**
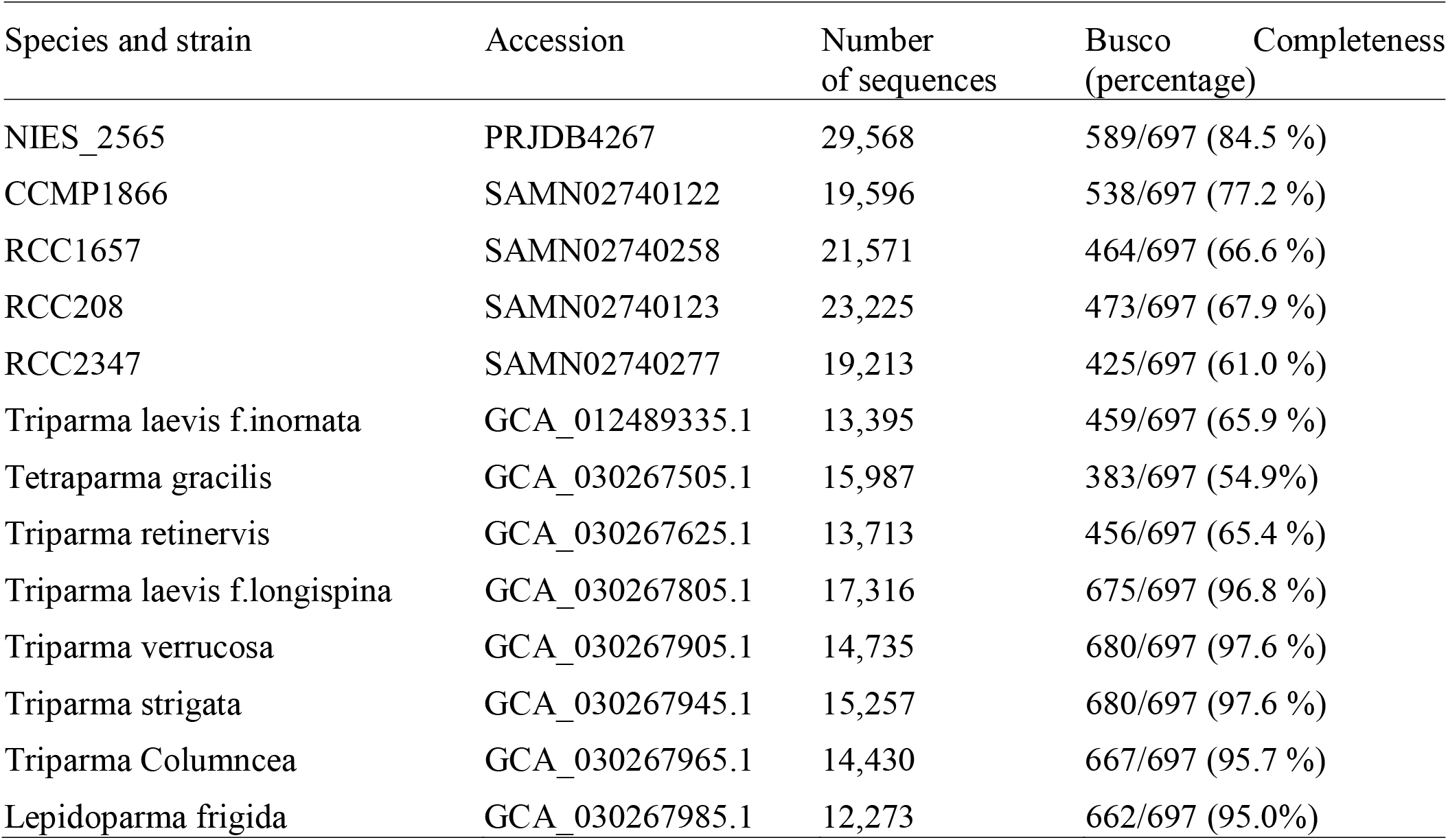
Comparison of assembly statistics and BUSCO completeness between the de novo transcriptomes and reference Parmales genomes.

**Table 2.** Coefficients of determination (*R*^2^) between S-type and F-type marker abundances in metagenomic and metatranscriptomic datasets.

| MAG | Dataset | Correlation ($R^2$) |
| --- | --- | --- |
| ARC-MAG | Metagenome | 0.5875 |
| ARC-MAG | Metatranscriptome | 0.3721 |
| PON-MAG | Metagenome | 0.8143 |
| PON-MAG | Metatranscriptome | 0.5834 |

Mapping *Tara* Oceans metatranscriptomic reads to the two MAGs revealed that all of the S-type and F-type marker genes encoded in the MAGs were transcriptionally active at least in one sample. Hierarchical clustering of expression profiles showed that S-type and F-type markers formed distinct clusters within each MAG (**Fig. 3**), indicating that the two marker sets are independently regulated.

**Figure 3.**
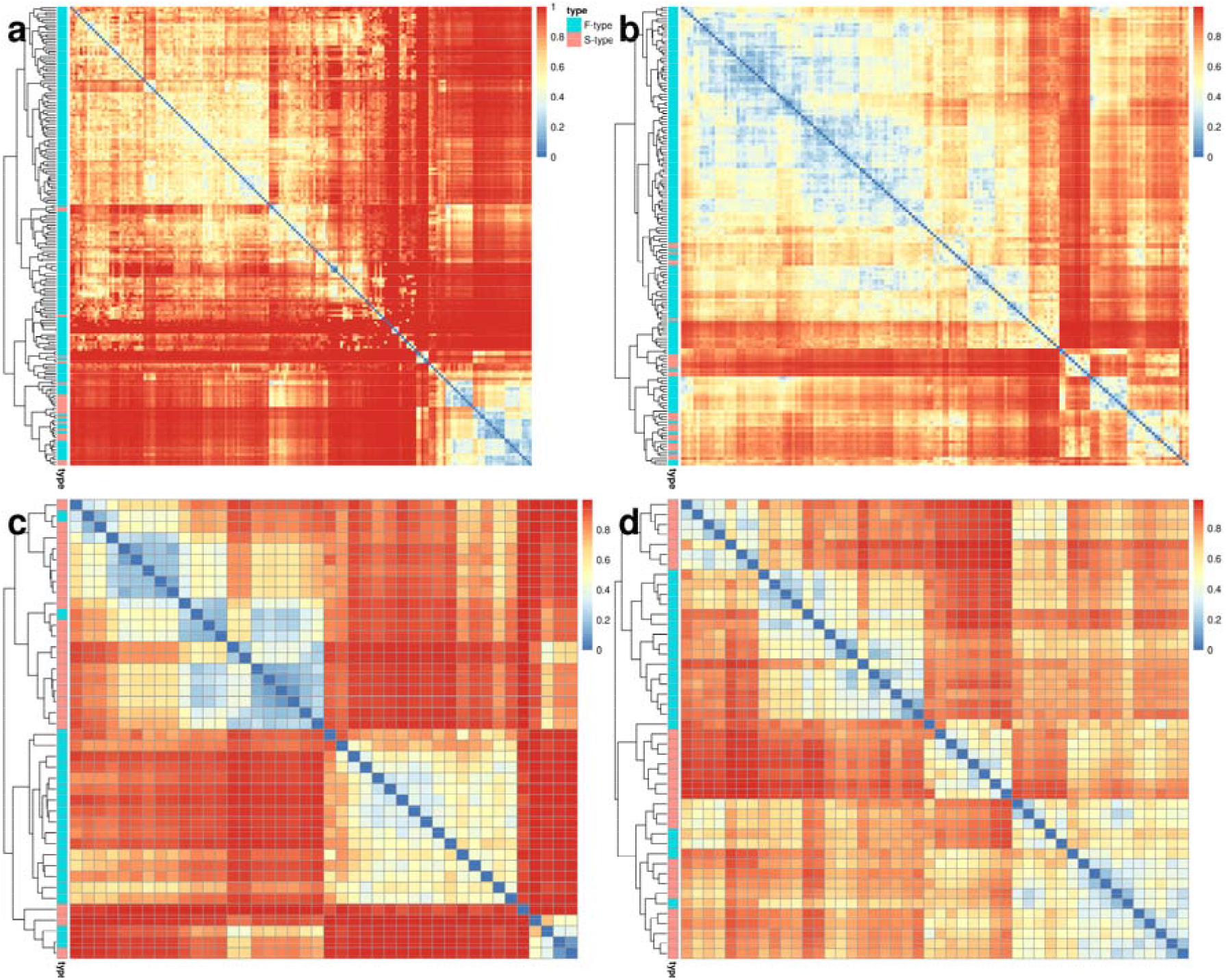
Expression profiles of S-marker and F-marker genes in two parmalean MAGs. Heatmaps displaying the similarity of expression patterns among S- and F-markers. The Bray-Curtis dissimilarity among marker genes was calculated based on the read counts of genes among samples. The color scale indicates the dissimilarity of expression patterns across orthologs. The annotation bars on the top and left indicate the morphological types: green for F-typ and red for S-type. (a, b) show the clustering of all marker genes for the Arctic MAG (a) and Pacific Ocean North MAG (b). (c, d) show the clustering of the 50 most highly expressed F-markers and all S-markers for the Arctic MAG (c) and Pacific Ocean North MAG (d).

Spatially, metagenomic distributions were synchronized between markers, whereas transcriptomic patterns diverged (**Fig. 4**). ARC-MAG transcription was restricted to polar regions, with F-type markers notably upregulated in the Atlantic inflow at the DCM layer (**Fig. 4d**). Conversely, PON-MAG transcripts were concentrated in low-to-middle latitudes, showing higher abundance in the DCM (Fig. 4h) than in surface waters (Fig. 4g). While F-type transcripts remained prevalent, S-type expression prominently increased within the DCM, especially in the Eastern Pacific.

**Figure 4.**
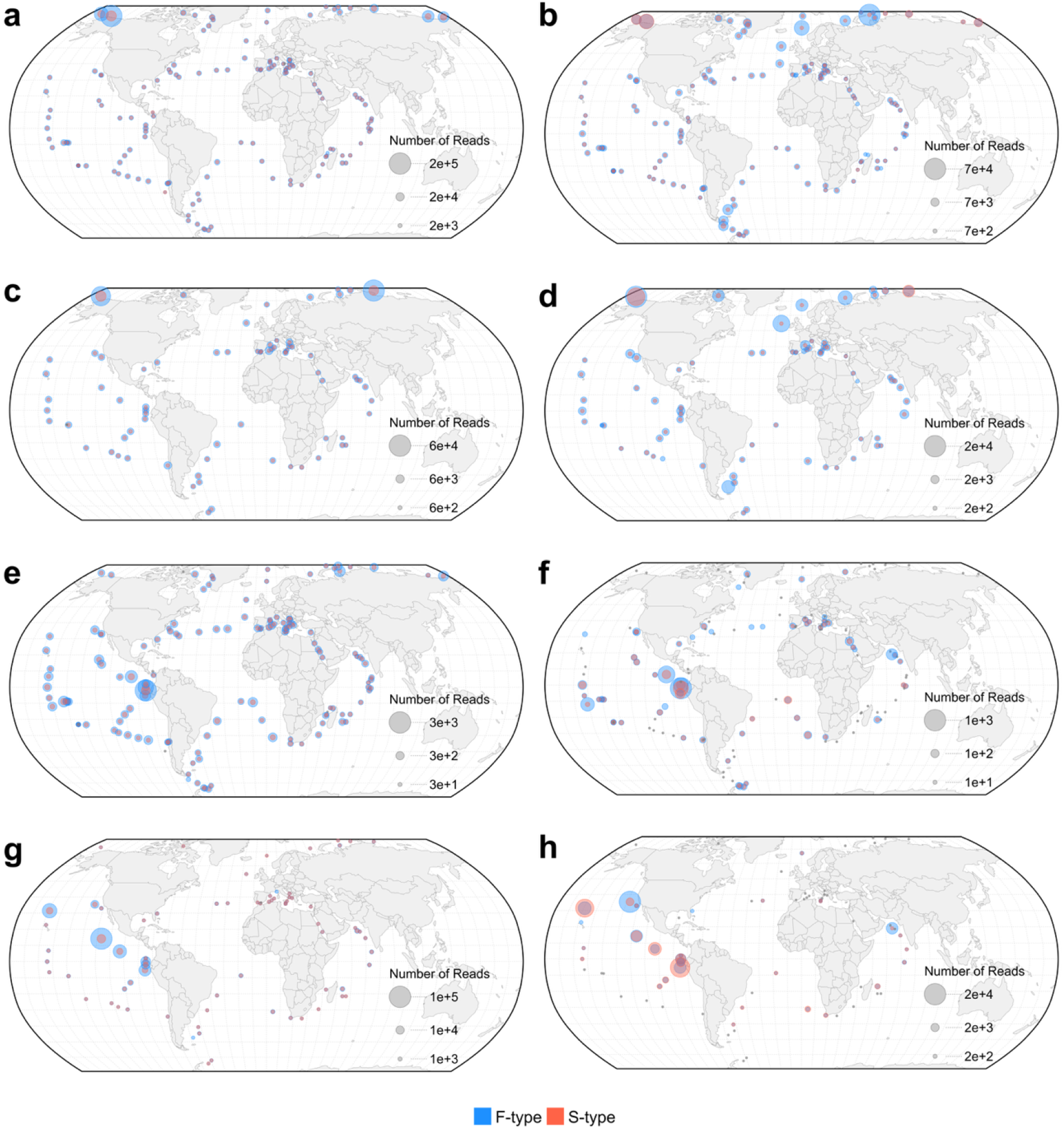
Global distribution and expression patterns of S- and F-type marker genes for two parmalean MAGs. The maps show read counts from metagenomes (left column: a, c, e, g) and metatranscriptomes (right column: b, d, f, h). Bubble size is proportional to the number of reads, and color indicates the marker type (blue: F-type; red: S-type). Sites with read counts < 10 are shown as dots. Samples are organized by MAG and depth layer: (a, b) ARC-MAG in surface waters; (c, d) ARC-MAG in the Deep Chlorophyll Maximum (DCM) layer; (e, f) PON-MAG in surface waters; (g, h) PON-MAG in the DCM layer.

### Distinct scaling regimes of morphotype-specific gene expression

The relationship between total gene abundance and S-type transcript abundance revealed two distinct scaling regimes (**Fig. 5**). In the high-expression regime (Group 1), S-type transcripts scaled near-linearly with gene abundance (slopes: 1.18 ± 0.12 for ARC-MAG, 0.90 ± 0.08 for PON-MAG; R_2_ = 0.82 and 0.86, respectively), indicating silicification gene expression proportional to relative genome abundance within the plankton community. In the basal regime (Group 2), scaling was markedly weaker (slopes: 0.54 ± 0.07 and 0.28 ± 0.07; R_2_ = 0.24 and 0.08), with S-type transcript levels remaining low and stochastic regardless of gene abundance. In contrast, F-type transcript abundance scaled uniformly with gene abundance without exhibiting such regime separation (slopes: 0.80 ± 0.09 and 0.87 ± 0.06; R_2_ = 0.28 and 0.54). These results suggest that S-type gene expression requires specific environmental triggers that lead to a higher population density, whereas F-type expression is constitutively maintained.

**Figure 5.**
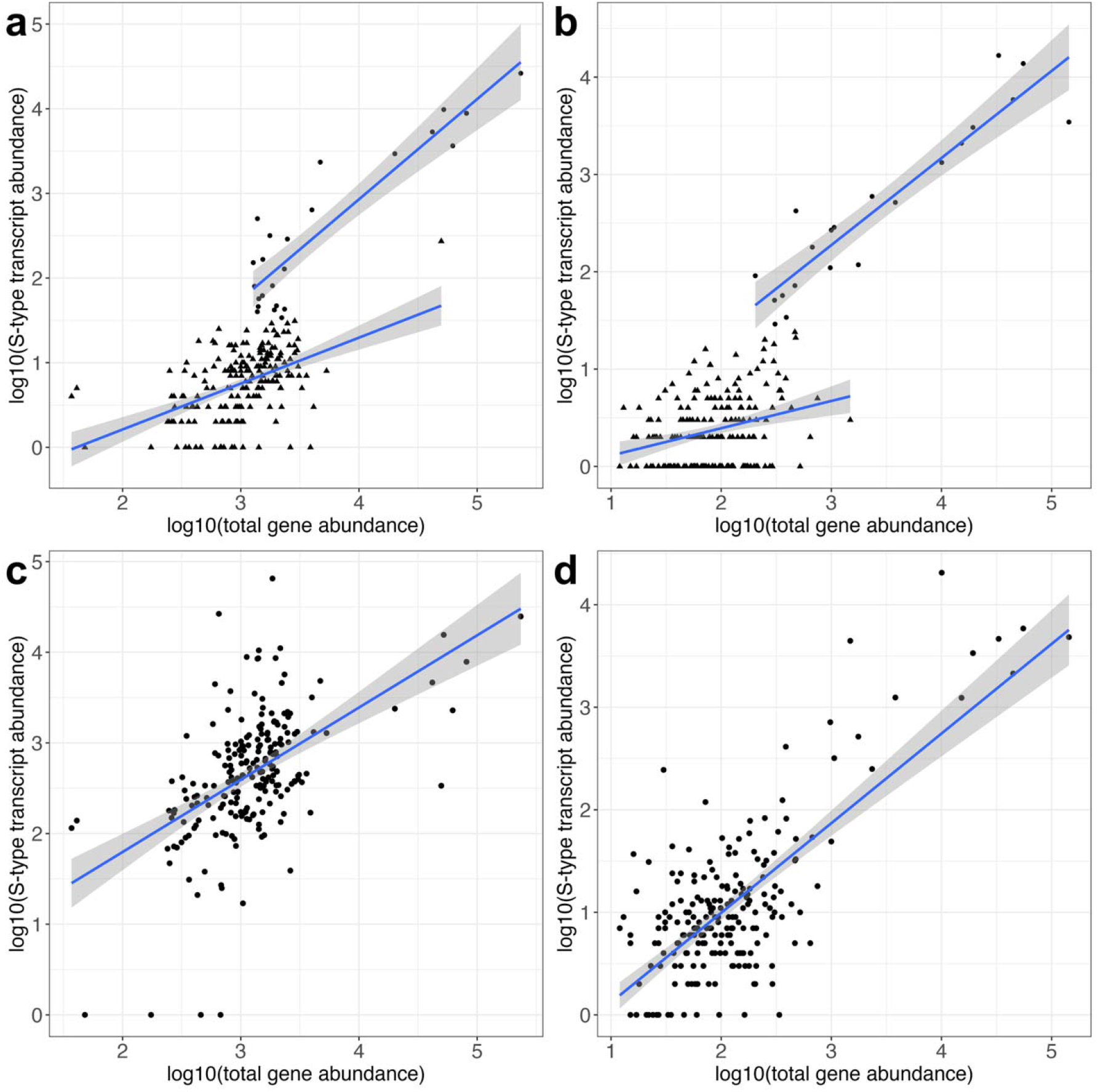
Scaling relationship between total gene abundance and morphotype-specific transcript abundance in ARC-MAGs and PON-MAGs. Log-log scatter plots showing the relationship between log10-transformed total gene abundance (metaG) and marker transcript abundance (metaT). (a, b) S-type markers for ARC-MAG and PON-MAG, respectively. Data points are divided into two groups: a manually defined subset showing a proportional response (Group 1, circles) and the remaining population (Group 2, triangles). (c, d) F-type markers for ARC-MAG and PON-MAG, respectively, showing uniform scaling without distinct regimes. Blue lines indicate linear regression fits with 95% confidence intervals (grey shading).

### Environmental drivers of morphology-specific gene expression

We further explored the environmental parameters that may affect expression profiles of morphotypes marker genes. GAMs revealed contrasting environmental dependencies between the two marker types (**Fig. 6, Fig. S3**). Notably, S-type models explained substantially more deviance (48.8% for ARC-MAG, 26.0% for PON-MAG) than F-type models (19.7% and 5.8%), indicating that S-type expression is more tightly coupled to abiotic variables. In the ARC-MAG (n=146), S-type expression was significantly associated with PO_4_, Si, temperature, salinity, and total carbon, whereas F-type expression responded primarily to temperature and NO_2_+NO_3_. In the PON-MAG (n=142), temperature was the dominant driver for both types, with PO_4_ additionally influencing S-type and NO_2_+NO_3_ influencing F-type expression. Sampling depth was not significant in any model.

**Figure 6.**
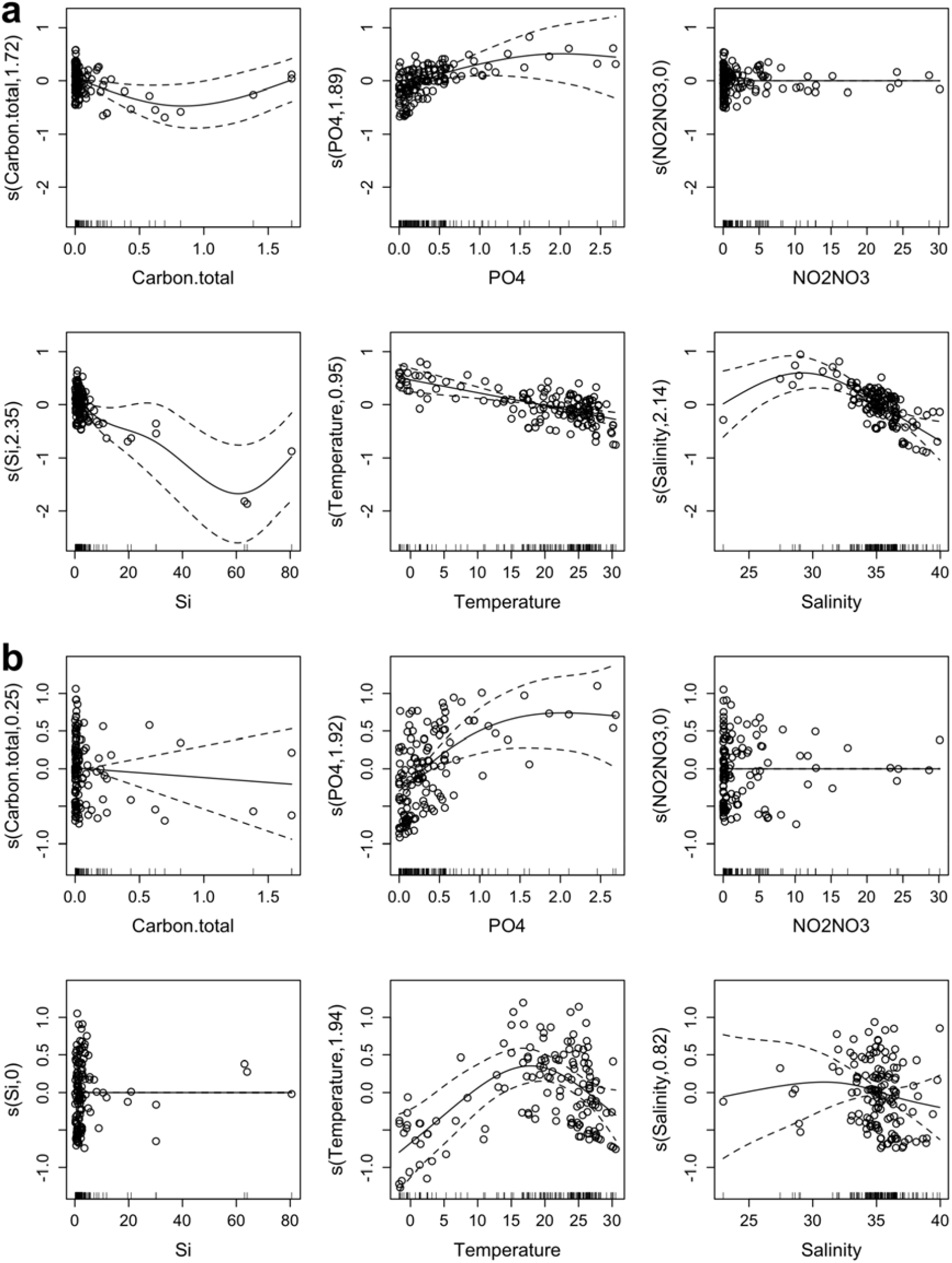
Generalized Additive Model (GAM) regression analysis of S-type specific expression variations based on environmental factors. Panels show partial effects of environmental variables on expression indices for the ARC-MAG dataset (n=146; a) and the PON-MAG dataset (n=142; b). The y-axis represents the partial effect (centered at zero) of each environmental variable on the expression index. Solid lines indicate the fitted smoothing curves, dashed lines represent 95% confidence intervals, and open circles show the partial residuals.

## Discussion

In this study, we detected the expression of marker genes for both S- and F-type forms from individual parmalean MAGs. While Ban et al. (2023) previously demonstrated that parmaleans (S-type) possess the genomic repertoire for these distinctive functions (Ban et al., 2023), our findings indicate, for the first time, that environmental parmalean populations (at the species level) manifest gene expression for both morphotypes. Previously, the hypothesis that these two forms belong to the same species was primarily inferred from molecular phylogenies (Ichinomiya et al., 2016). Our results support the view that these dual genetic traits are functional in actual marine environments, at least within the lineages represented by the two analyzed MAGs.

The global distribution patterns of the two MAGs align well with the biogeography of parmalean clades elucidated by Ban et al. (2024): ARC-MAG was predominantly found in polar and subarctic regions, mirroring Clade I (Lepidoparmales), while PON-MAG was detected in tropical and temperate regions, corresponding to the warmer-water populations within the ubiquitous Clade IV.

A notable observation was the contrasting distribution patterns between S-type and F-type marker genes at the transcriptomic level, despite their stable genomic ratios across all sites. This genomic stability indicates that the geographically distinct expression patterns are driven by differential transcriptional activity rather than differences in intra-species genetic variations. This finding offers a coherent explanation for the long-standing discrepancy between the global ubiquity reported in Amplicon Sequence Variant (ASV)-based surveys and the restricted range of silicified strain isolations: populations in warmer waters are likely represented primarily by the flagellated form.

The restriction of the S-type phase appears to be governed by environmental parameters. Our scaling analysis revealed two distinct regimes of S-type gene expression (**Fig. 5**), which, together with the environmental modeling (**Fig. 6**), suggests that conditions favorable for biomass increase trigger active silicification. In short, S-type cells appear to increase only under conditions favorable for growth, while F-type cells appear to tolerate various conditions but their response to growth-favoring condition is not as sharp as S-type. This interpretation aligns with culture experiments showing that the S-type is cold-adapted with limited growth above 15°C (Ichinomiya et al., 2013), whereas the F-type exhibits broader thermal tolerance (Stawiarski et al., 2016). Thus, the “ubiquitous” distribution of Clade IV in ASV studies likely reflects the capacity of the F-type to inhabit warmer waters, while the S-type is confined to niches satisfying specific growth conditions.

This niche separation is illustrated by the PON-MAG in tropical regions, where S-type expression was substantially lower in surface waters than in the DCM, consistent with microscopic observations of silicified parmaleans in deep photic layers (Fujita & Jordan, 2017). The historical reports of parmalean “absence” in tropical surface waters likely reflect suppressed silicification below detection limits. Conversely, F-type markers were robustly detected in these surface layers, suggesting that the flagellated form inhabits this warm niche, remaining cryptic to morphological observation. The presence of the F-type in these typically oligotrophic surface waters may reflect differences in trophic strategies. Ban et al. (2023) hypothesized dimorphic trophic modes: primarily autotrophic for the S-type and phago-mixotrophic for the F-type (Ban et al., 2023). Our data are compatible with this hypothesis, as S-type markers prevailed in the nutrient-rich DCM whereas F-type markers dominated in nutrient-depleted surface waters where bacterial grazing could sustain a mixotroph. Nevertheless, since the flagellated form is hypothesized to function as a gamete, its surface presence could partly reflect life-cycle transitions.

Our GAM analysis provides statistical support for these divergent strategies. S-type models demonstrated substantially higher explanatory power than F-type models, indicating that S-type expression is strongly coupled to abiotic variables while F-type expression is largely decoupled from measured physicochemical parameters. This contrast reinforces the hypothesis of distinct trophic modes: strict autotrophy would be directly regulated by inorganic nutrients, whereas a mixotrophic F-type driven by prey availability would not be captured by standard environmental metadata. Among the specific drivers, S-type expression showed consistent positive relationships with PO_4_ in both MAGs. For the ARC-MAG, the additional association with lower salinity suggests an affinity for river-influenced, nutrient-rich Arctic waters, while the negative relationship with Si may reflect rapid drawdown by actively silicifying populations.

Building on these contrasting environmental dependencies, we propose a threshold-based model for the morphological switch in parmaleans. Our scaling analysis revealed that S-type accumulation consistently occurred above a minimum abundance threshold, suggesting that the transition from flagellated to silicified phases is not gradual but requires conditions sufficient to sustain stable growth. Below this threshold—as observed in oligotrophic surface waters—the community appears to persist primarily in the F-type form, whose mixotrophic capacity enables survival in nutrient-poor environments where strict autotrophy would be untenable. Once nutrient and other conditions reach a certain level that sustain S-type growth, the morphological switch is triggered and both phases accumulate within the community. This model reconciles the cryptic, F-type-dominated populations in warm oligotrophic waters with the conspicuous blooms of silicified cells reported in nutrient-rich regions (11).

These insights must be interpreted with caution. The low abundance of Parmales [<0.2% of the community (Ichinomiya et al., 2016; Kuwata et al., 2018)] limited the number of mapped reads, and missing environmental metadata necessitated sample exclusions that may have introduced bias. To validate the life cycle model, it is necessary to move beyond static distributional snapshots. While culture-based direct observation of S–F transitions remain the ultimate goal, field-based temporal dynamics data would offer an important alternative. Priority should be given to high-abundance regions absent from our dataset, such as the Western North Pacific (Oyashio) and the east coast of Australia, where Clade II blooms are known to occur (Ban et al., 2024). Integrating targeted environmental profiling will be an effective approach to elucidate the environmental drivers of morphological switching in this unique lineage.

## Acknowledgement

We thank the *Tara* Oceans consortium and the people and sponsors who supported the *Tara* Oceans Expedition (http://www.embl.de/tara-oceans/) for making the data accessible. This is contribution number XXX of the *Tara* Oceans Expedition 2009–2013. Computational time was provided by the Supercomputer System, Institute for Chemical Research, Kyoto University.

