## Supplementaly Figures for "Metatranscriptome data support the existence of two distinct morphotypes in a single parmalean species in natural environments"

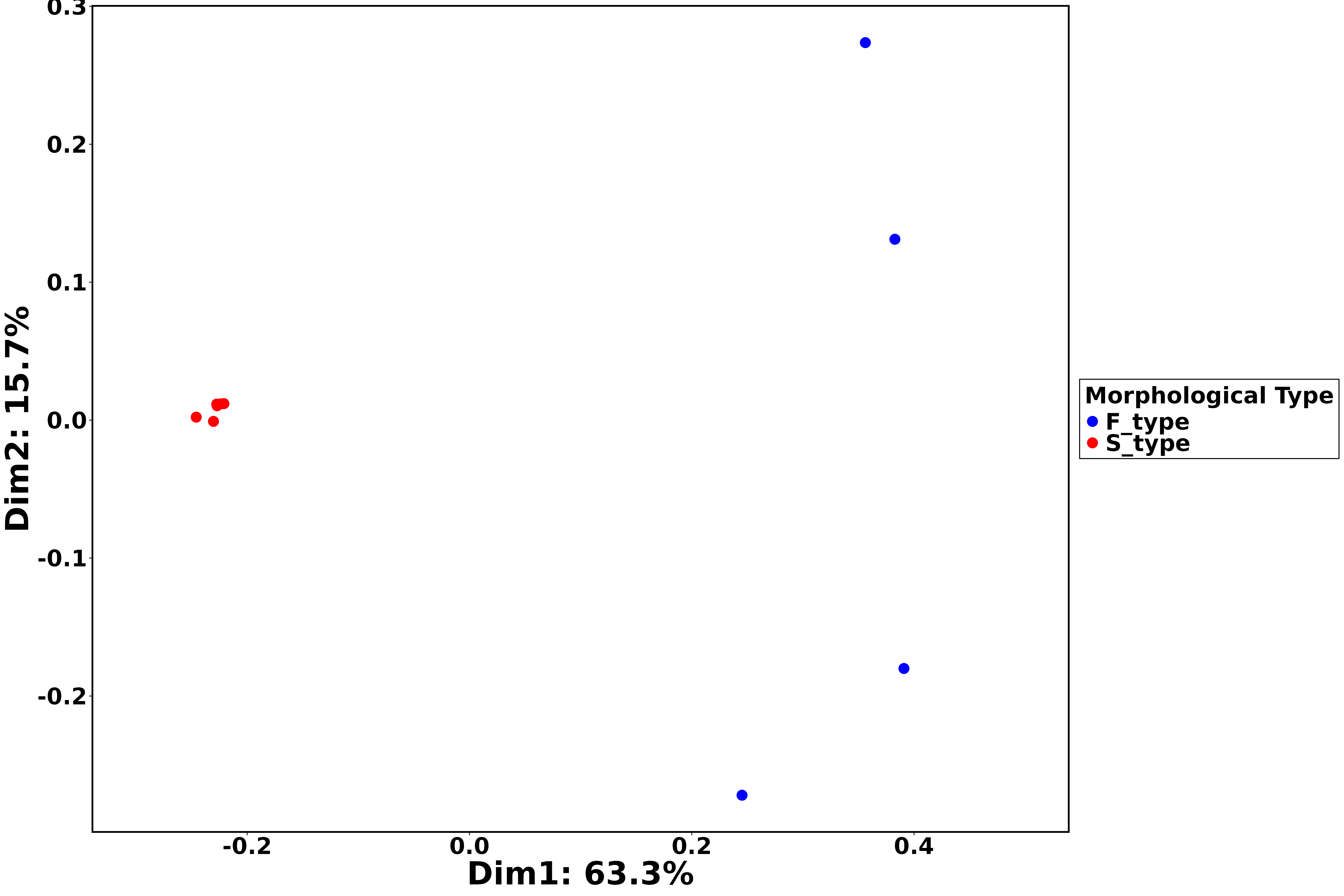

**Figure S1. PCoA plot of gene expression profiles.** The plot is based on Bray-Curtis dissimilarity calculated from read counts mapped to orthologous genes across 10 RNA-seq samples. Each point represents an individual RNA-seq sample. Blue points indicate the four F-type strains, while red points indicate the six S-type samples (including both silicate-rich and silicate-deficient conditions).

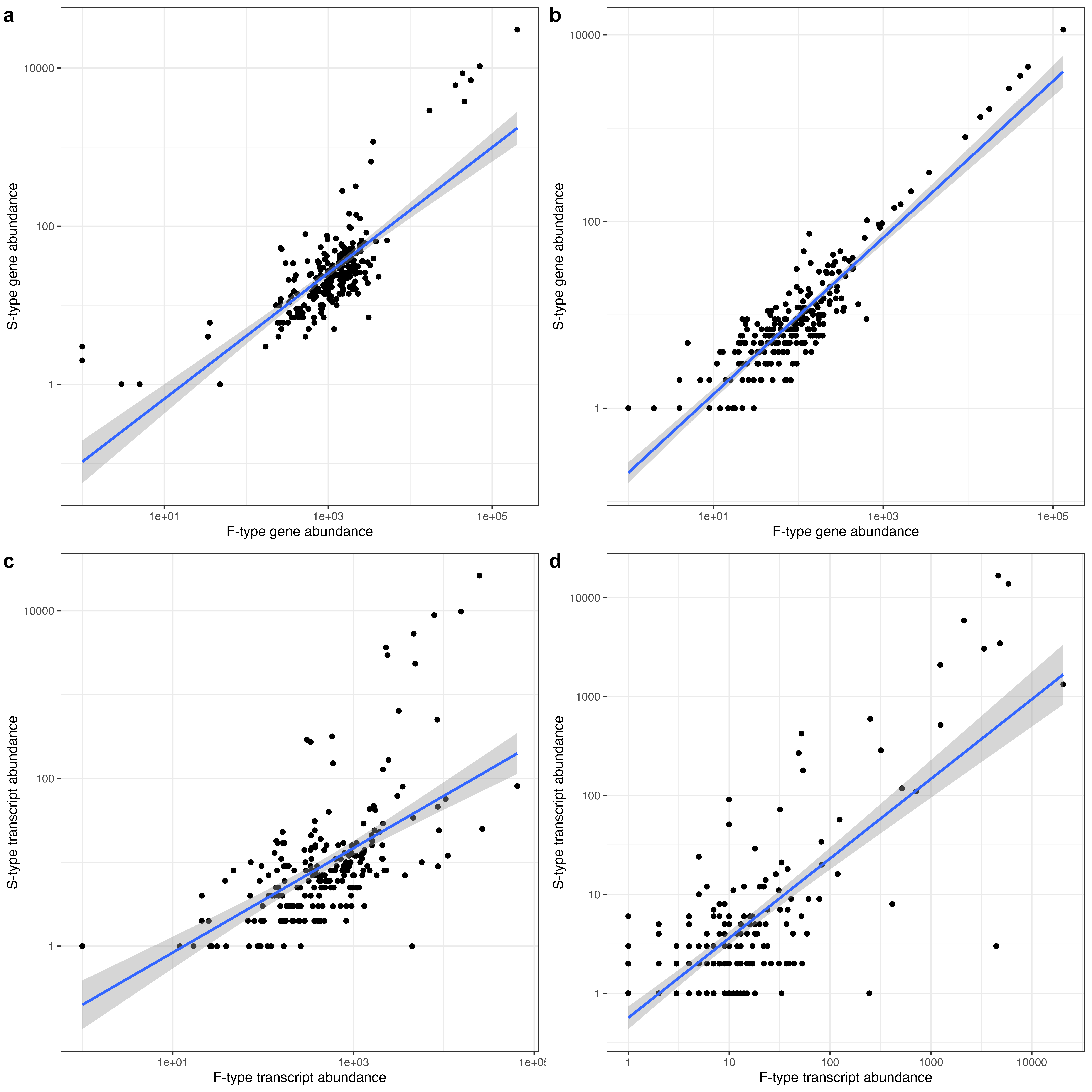

**Figure S2. Comparison of genomic and transcriptomic abundance ratios between S- and F-type markers.** Relationship between F-type and S-type marker abundances for ARC-MAG (a, c) and PON-MAG (b, d). (a, b) Metagenomic abundance showing consistent gene ratios across global sites. (c, d) Metatranscriptomic abundance showing variable transcript ratios, indicating differential expression. Each point represents a sampling site. Blue lines indicate linear regression fits with 95% confidence intervals (gray shading). Both axes are log-scaled.

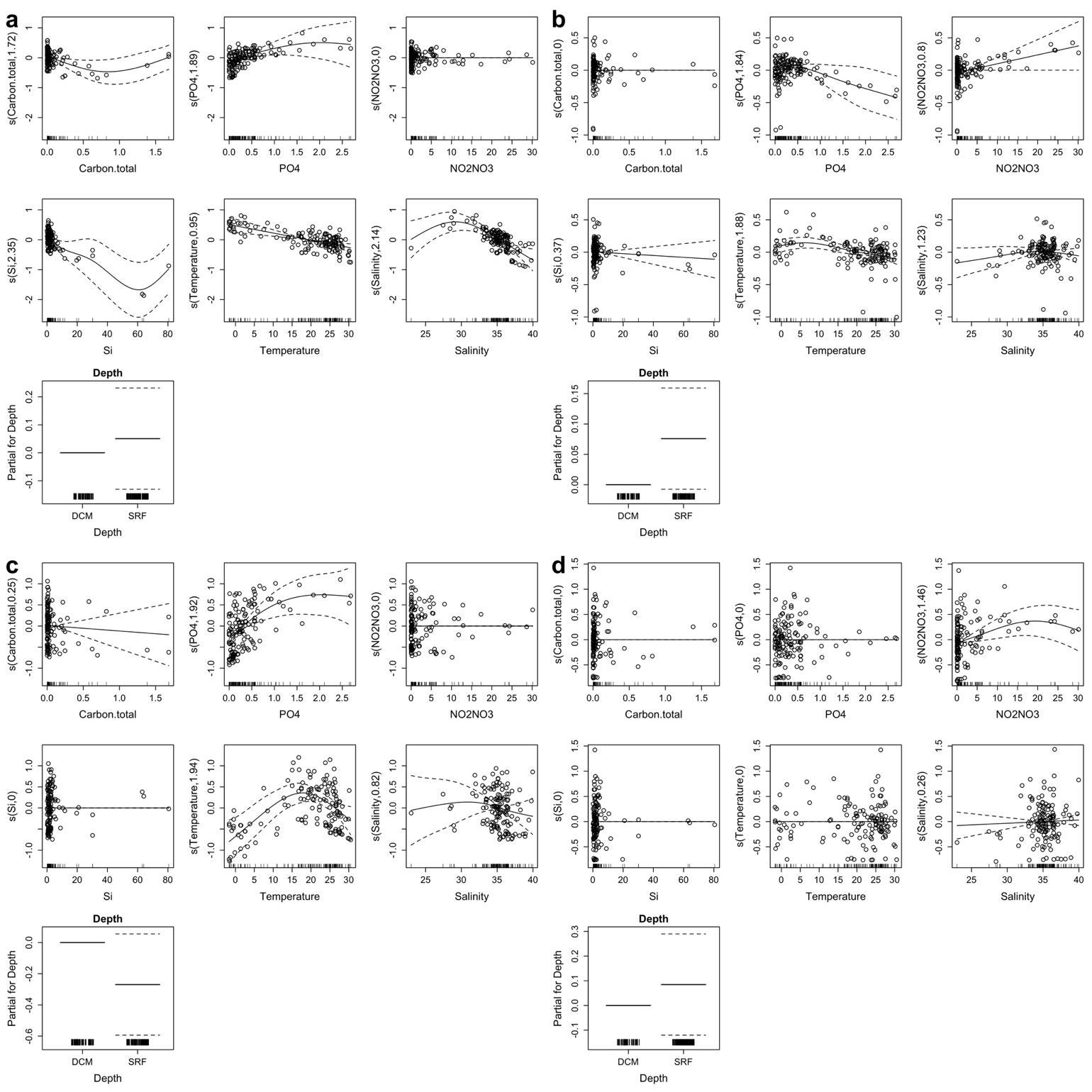

**Figure S3. Generalized Additive Model (GAM) regression analysis of morphotype-specific expression variations based on environmental factors.** Panels show partial effects of environmental variables on expression indices for the ARC-MAG dataset (n=146; a, b) and the PON-MAG dataset (n=142; c, d). Specifically, panels (a, c) correspond to S-type marker genes, while panels (b, d) correspond to F-type marker genes. The y-axis represents the partial effect (centered at zero) of each environmental variable on the expression index. Solid lines indicate the fitted smoothing curves, dashed lines represent 95% confidence intervals, and open circles show the partial residuals.
